# Gain-of-function mutation in *Gli3* causes ventricular septal defects

**DOI:** 10.1101/2020.02.10.942144

**Authors:** Antonia Wiegering, Paniz Adibi, Ulrich Rüther, Christoph Gerhardt

**Affiliations:** Institute for Animal Developmental and Molecular Biology, Heinrich-Heine University Düsseldorf, 40225 Düsseldorf, Germany; Department of Pediatric Cardiology, Justus Liebig University, 35390 Gießen, Germany

**Keywords:** congenital heart defects, Hedgehog signalling, PDGFRα signalling, primary cilia, cell proliferation

## Abstract

Ventricular septal defects (VSDs) are developmental disorders, characterised by a gap in the septum between the right and the left ventricle, that lead to life-threatening heart defects. At present, the only curative treatment of VSDs is surgical closure. Since these surgeries comprise several severe risks, the development of alternative therapies against VSDs is urgently needed. To develop such therapies, the current knowledge of the molecular factors and mechanisms underlying VSDs has to be increased. Based on our previous data, we analysed the relevance of the HH signalling pathway mediator GLI3 in ventricular septum (VS) formation. GLI3 functions as both a transcriptional activator (GLI3-A) and repressor (GLI3-R). By analysing two different mouse *Gli3* mutants, we revealed that the lack of GLI3-A with simultaneous presence of GLI3-R impairs cilia-mediated PDGFRα signalling causing reduced cell proliferation and in consequence the development of VSDs. Moreover, we showed that the rescue of PDGFRα signalling restores cell proliferation. Since VSDs are also appear in humans with comparable gain-of-function mutations in *GLI3*, our findings propose activators of PDGFRα signalling as potential agents against the development of VSDs.

**SUMMARY:** The article reports how a gain-of-function mutation of *Gli3* causes ventricular septal defects and paves the way for therapies tackling these congenital heart defects.

## INTRODUCTION

In the vertebrate heart, the VS localises between the right and the left ventricle and is crucial for the separation of oxygenated and deoxygenated blood. Any opening in the VS represents a VSD which results in the mixture of oxygenated and deoxygenated blood. Larger VSDs often lead to an increased blood flow towards the lung and the left ventricle (Brickner et al., 2000) and hence to the development of left ventricular hypertrophy and pulmonary edema and dilatation (Ferreira et al., 2009; Selicorni et al., 2009). In a former study, we revealed that cardiac primary cilia are essential for VS formation (Gerhardt et al., 2013). Primary cilia are tiny cytoplasmic protrusions consisting of the basal body (BB), the axoneme, the transition zone (TZ) and the ciliary membrane (Reiter and Leroux, 2017; Satir et al., 2010). The BB is derived from the mother centriole and gives rise to the ciliary microtubule-based scaffold, the axoneme. The axoneme comprises nine doublet microtubules arranged in a circle and is essential for the transport of proteins that are conveyed through the cilium. At the proximal part of the axoneme, the TZ governs ciliary protein import and export. On the surface, cilia are confined by the ciliary membrane. Primary cilia mediate various signalling pathways and one of these pathways is the Hedgehog (HH) signalling cascade (Anvarian et al., 2019; Satir et al., 2010; Wheway et al., 2018). HH signalling starts when the ligand HH [e.g. Sonic hedgehog (SHH)] binds to its receptor Patched 1 (PTCH1), which is located in the ciliary membrane. This binding event is supported by Low-density lipoprotein receptor related protein 2 (LRP2) (Christ et al., 2012). The resulting HH/PTCH1 complex leaves the cilium and Smoothened (SMO) is able to access the ciliary membrane (Corbit et al., 2005; Rohatgi et al., 2007). There, SMO separates the full-length Glioma-Associated Oncogene Family Zinc Finger 2 (GLI2) and Glioma-Associated Oncogene Family Zinc Finger 3 (GLI3) proteins from Suppressor of Fused (SUFU) by initiating a yet unknown mechanism and converts them into transcriptional activators (GLI2-A and GLI3-A) (Chen et al., 2009; Humke et al., 2010). This conversion is promoted by the proteins Broad-Minded (BROMI; also referred to as TBC1D32), Ellis Van Creveld 1 (EVC1) and Ellis Van Creveld 2 (EVC2) (Blair et al., 2011; Caparrós-Martín et al., 2013; Dorn et al., 2012; Ko et al., 2010; Ruiz-Perez et al., 2007; Valencia et al., 2009; Yang et al., 2012). GLI2-A and GLI3-A exit the cilium, enter the nucleus and induce HH target gene expression (e.g., the expression of *Gli1* or *Ptch1*). The intraciliary transport of PTCH1, SMO and GLI2 is implemented by the Intraflagellar transport proteins 25 and 27 (IFT25 and IFT27) (Eguether et al., 2014; Keady et al., 2012; Yang et al., 2015). Thus, the loss of either IFT25 or IFT27 results in a decreased expression of HH target genes (Eguether et al., 2014; Keady et al., 2012). The lack of HH leads to the continued presence of PTCH1 within the ciliary membrane. In turn, SMO remains outside the cilium leading to a proteolytic processing of the full-length GLI2 and GLI3 proteins (GLI2-FL and GLI3-FL) into transcriptional repressors (GLI2-R and GLI3-R) (Gerhardt et al., 2015; Gerhardt et al., 2016b; Wang et al., 2000). GLI2 and GLI3 processing begins with the phosphorylation of GLI2 and GLI3 by protein kinase A (PKA), Caseinkinase1 (CK1) and Glycogensynthasekinase3-β (GSK3-β) (Pan et al., 2006; Tuson et al., 2011; Wang and Li, 2006). In addition, the proteins Kinesin family member 7 (KIF7) and Fuzzy (FUZ) participate in this processing event (Brooks and Wallingford, 2012; Dai et al., 2011; Du Toit, 2014; Gray et al., 2009; He et al., 2014; Heydeck et al., 2009; Pedersen and Akhmanova, 2014; Zilber et al., 2013). Finally, GLI processing is implemented by the cilia-regulated proteasome (Gerhardt et al., 2015; Gerhardt et al., 2016b).

It is assumed that HH signalling controls proper VS development as the loss or the truncation of proteins that positively regulate HH signalling causes the occurrence of VSDs. For example, defects in ventricular septal formation are observed in *Shh, Lrp2, Sufu, Bromi, Ift25* and *Ift27* mutant mouse embryos (Eguether et al., 2014; Keady et al., 2012; Li et al., 2015; Washington Smoak et al., 2005), while mutations in the *EVC* gene lead to VSDs in humans (Liu et al., 2018). But, remarkably, mutations in *Gsk3-β, Kif7* and *Fuz* encoding negative regulators of HH signalling also result in the occurrence of VSDs in mice (Coles and Ackerman, 2013; Kerkela et al., 2008; Li et al., 2015) indicating that the inhibition of HH target gene expression might be important for proper VS formation. Interestingly, GSK3-β, KIF7 and FUZ negatively regulate HH signalling by supporting proteolytic processing of GLI2 and GLI3. The general view is that GLI3-R functions as the main transcriptional repressor of the HH pathway (Wang et al., 2000). Previously, we reported that the loss of the ciliary protein RPGRIP1L results in the occurrence of VSDs in mice and suggested that these VSDs are caused by an impaired GLI3 processing (Gerhardt et al., 2013). However, the lack of GLI3 in mice (*Gli3*^*XtJ/XtJ*^ mice) does not result in the development of VSDs (Johnson, 1967).

To shed light on the role of GLI3 in VS development, we analysed *Gli3*^*XtJ/XtJ*^ mice and mouse embryos which produce GLI3-R but lack GLI3-A (*Gli3*^*Δ699/Δ699*^ embryos) (Böse et al., 2002; Hill et al., 2007; Johnson, 1967). In contrast to *Gli3*^*XtJ/XtJ*^ embryos, *Gli3*^*Δ699/Δ699*^ embryos displayed VSDs. Investigating the molecular causes underpinning these VSDs, we revealed reduced HH signalling in *Gli3*^*Δ699/Δ699*^ embryonic hearts. The decreased HH signalling led to diminished cilia-mediated PDGFRα signalling which, in turn, caused a reduced proliferation in those parts of the cardiac walls that contribute to the formation of the VS. *In vitro* studies with PDGFRα pathway activators suggest that a restoration of PDGFRα signalling might rescue VS formation in *Gli3*^*Δ699/Δ699*^ mouse embryos.

## RESULTS AND DISCUSSION

### *Gli3*^*Δ699/Δ699*^ mouse embryos develop VSDs and exhibit reduced cardiac HH signalling

It is known that HH signalling participates in the formation of the VS but the role of the HH mediator GLI3 in this process is unclear (Gerhardt et al., 2013; Wiegering et al., 2017). While we could not detect VSDs in any analysed *Gli3*^*XtJ/XtJ*^ mouse embryo, 100% of the examined *Gli3*^*Δ699/Δ699*^ embryos (14 out of 14) displayed VSDs (Fig. 1A). This finding demonstrates that GLI3 is dispensable for VS formation but that the presence of GLI3-R alone causes VSDs. To analyse HH signalling in these mutants, we quantified the expression of the HH target genes *Gli1* and *Ptch1*. The expression of both genes was increased in the hearts of *Gli3*^*XtJ/XtJ*^ embryos and decreased in the hearts of *Gli3*^*Δ699/Δ699*^ embryos (Fig. 1B, C). Importantly, the amount of GLI2-A and GLI2-R was equal in the hearts of both mutants (Fig. 1D). Our data demonstrate that HH signalling is needed for proper VS formation as we detected VSDs in mouse embryos which produce GLI3-R but lack GLI3-A (Fig. 1A) and hence display reduced HH signalling (Fig. 2B, C). But the question remains open why VSDs develop in several mouse mutants lacking negative regulators of HH signalling (Coles and Ackerman, 2013; Kerkela et al., 2008; Li et al., 2015). A possible answer to this question might be that KIF7 and FUZ are also involved in the transformation of GLI2-FL and GLI3-FL into GLI2-A and GLI3-A by yet unknown mechanisms (Cheung et al., 2009; Endoh-Yamagami et al., 2009; He et al., 2014; Heydeck et al., 2009; Pedersen and Akhmanova, 2014). Moreover, it is conceivable that VSDs occur in these mutants since GSK3-β, KIF7 and FUZ also play a role in other signalling cascades that regulate the development of the VS such as TGF-β, Notch and WNT signalling (Chapman et al., 2019; Choi et al., 2007; Choudhary et al., 2006; Donovan et al., 2002; Fischer et al., 2007; Greenway et al., 2009; High et al., 2009; High et al., 2007; Jiao et al., 2003; Kokubo et al., 2004; Li et al., 1997; Liu et al., 2004; Nakajima et al., 2004; Oda et al., 1997; Rochais et al., 2009; Sakata et al., 2002; Song et al., 2010; Touma et al., 2017; Wurdak et al., 2005). It was already shown that GSK3-β is involved in TGF-β, Notch and WNT signalling (Aberle et al., 1997; Espinosa et al., 2003; Foltz et al., 2002; Guo et al., 2008; Kim et al., 2009) and that FUZ takes part in WNT signalling (Zilber et al., 2013).

**Figure 1:**
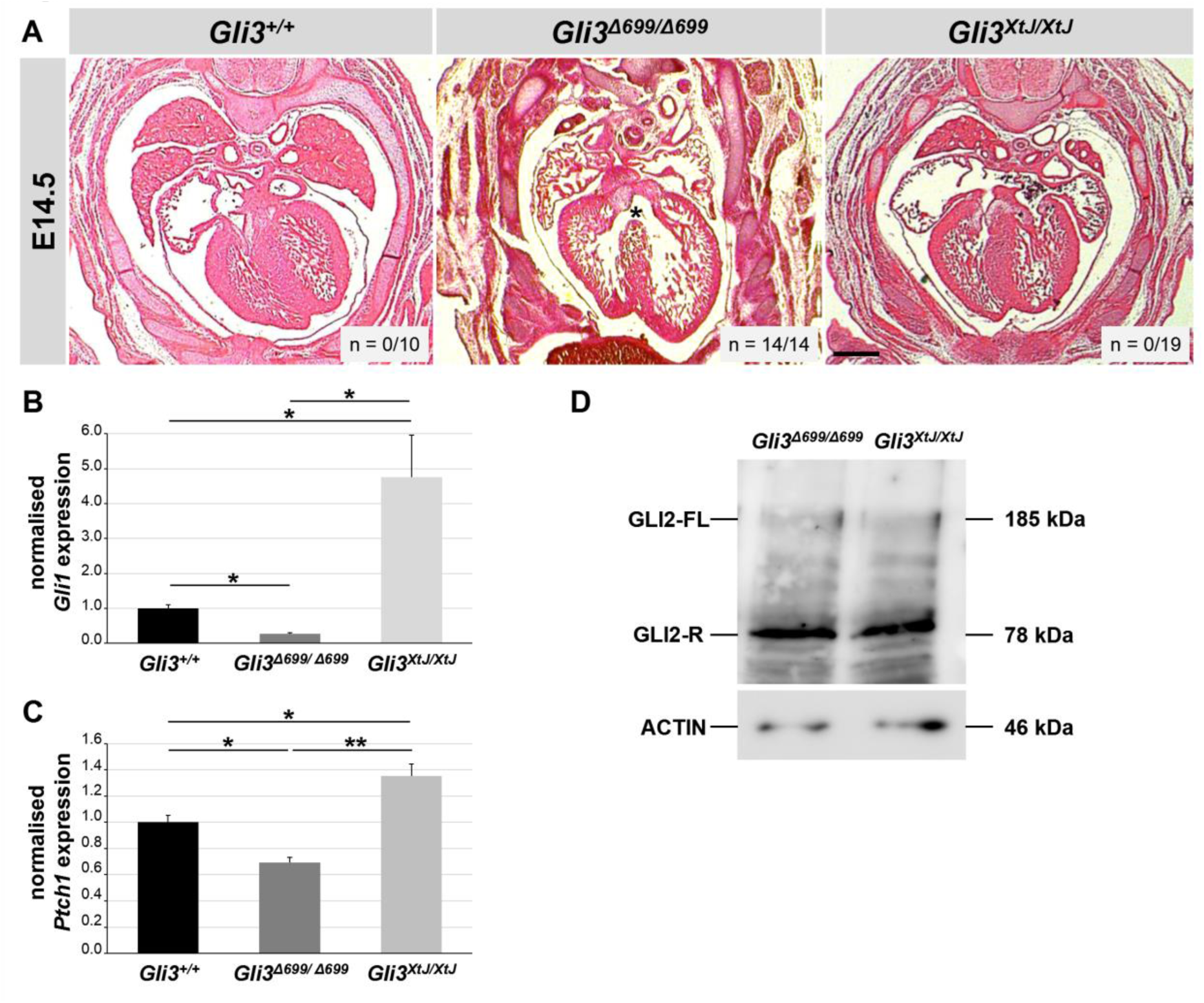
*Gli3*^*Δ699/Δ699*^ mutant embryos display VSDs and a decreased HH signalling. (A) HE staining of heart sections from WT, *Gli3*^*Δ699/Δ699*^ and *Gli3*^*XtJ/XtJ*^ embryos at E14.5. Asterisk indicates VSD. The scale bar represents a length of 500 µm. (B, C) Quantification of *Gli1* (B) and *Ptch1* (C) expression via qRT-PCR analysis. Lysates were obtained from WT, *Gli3*^*Δ699/Δ699*^ and *Gli3*^*XtJ/XtJ*^ (n = 3, respectively) mouse embryonic hearts at E12.5. (D) Western blot analysis of GLI2 with lysates obtained from *Gli3*^*Δ699/Δ699*^ and *Gli3*^*XtJ/XtJ*^ (n = 3, respectively) mouse embryonic hearts at E12.5. Actin serves as loading control.

**Figure 2:**
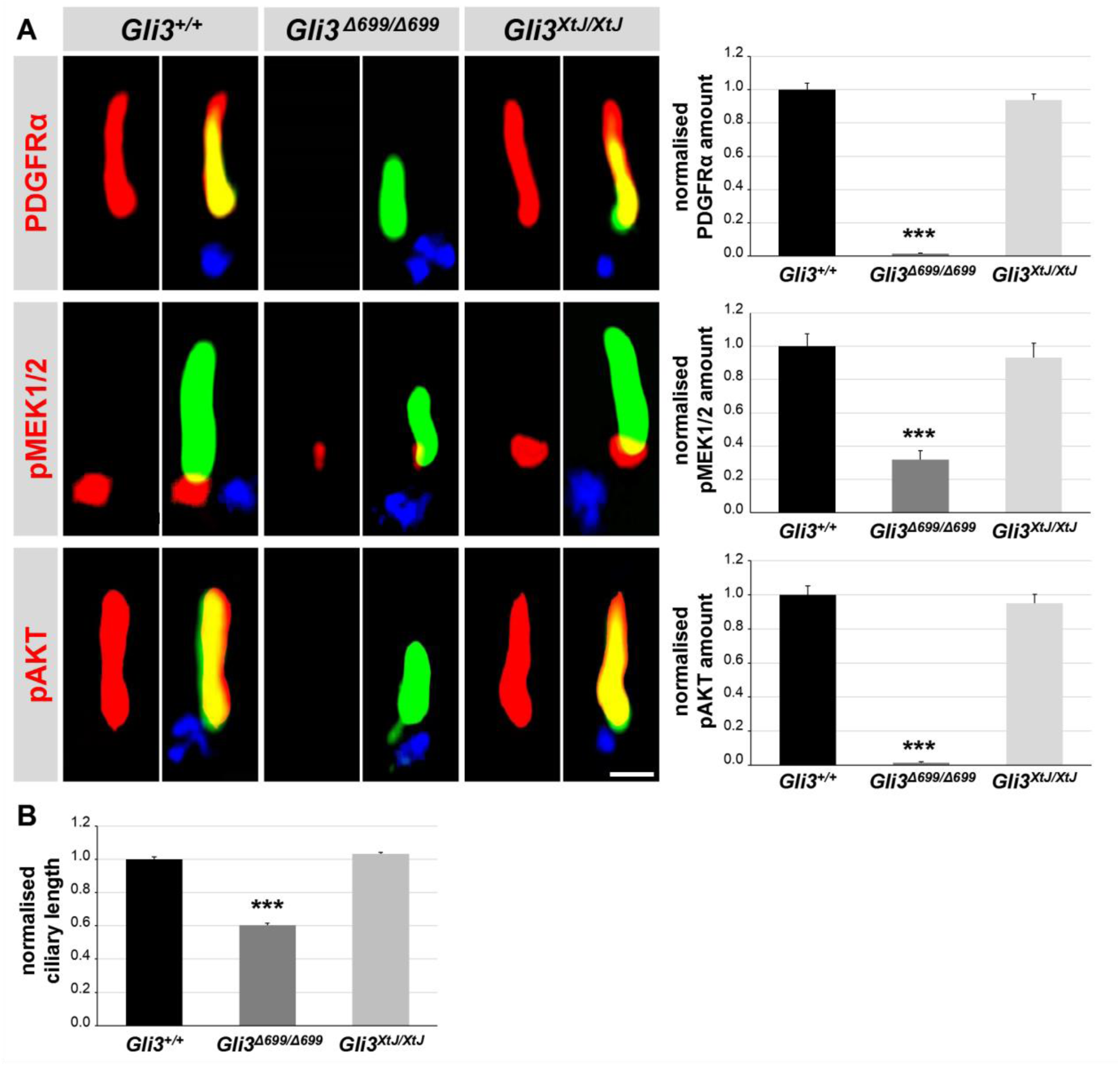
Reduced PDGFRα signalling and ciliary length in *Gli3*^*Δ699/Δ699*^ hearts. (A) Immunofluorescence on heart sections obtained from WT (n=12), *Gli3*^*Δ699/Δ699*^ (n=8) and *Gli3*^*XtJ/XtJ*^ (n=12) embryos at E12.5. The ciliary axoneme is stained in green by acetylated α-tubulin, the basal body is stained in blue by γ-tubulin. The scale bar represents a length of 0.5 µm. (B) Ciliary length measurement on mouse embryonic heart sections obtained from WT, *Gli3*^*Δ699/Δ699*^ and *Gli3*^*XtJ/XtJ*^ (n = 4, respectively) embryos at E12.5.

### PDGFRα signalling is reduced in ventricular cilia of *Gli3*^*Δ699/Δ699*^ embryonic hearts

We formerly showed that HH signalling controls the PDGFRα pathway in murine ventricles (Gerhardt et al., 2013). Based on this, we analysed different components of the PDGFRα pathway in the ventricles of *Gli3*^*XtJ/XtJ*^ and *Gli3*^*Δ699/Δ699*^ embryos (Fig. 2). It has been previously shown that PDGFRα signalling is mediated by primary cilia (Schneider et al., 2005). In wild-type ventricular cilia, PDGFRα is distributed along the entire cilium (Gerhardt et al., 2013). While it localised properly in ventricular cilia of *Gli3*^*XtJ/XtJ*^ embryos, PDGFRα was nearly vanished in ventricular cilia of *Gli3*^*Δ699/Δ699*^ embryos (Fig. 2A) reflecting that the presence of GLI3-R alone reduced the amount of ciliary PDGFRα. PDGFRα signalling starts, when its ligand PDGF-AA binds to PDGFRα which then becomes dimerised and phosphorylated. In turn, the phosphorylation of PDGFRα activates MEK1/2-ERK1/2 signalling as well as AKT signalling (Gerhardt et al., 2016a; Schneider et al., 2010; Schneider et al., 2005). Former reports depicted that activated (phosphorylated) MEK1/2 and activated (phosphorylated) AKT are also present in primary cilia (Clement et al., 2013; Schneider et al., 2010; Schneider et al., 2005; Struchtrup et al., 2018). For this reason, we next investigated whether pMEK1/2 and pAKT can be found in ventricular cilia of wild-type embryos. We detected pMEK1/2 at the transition zone of ventricular cilia and pAKT dispersed along ventricular cilia (Fig. 2A). Their ciliary amount was unaltered in *Gli3*^*XtJ/XtJ*^ embryonic ventricles and decreased in ventricular cilia of *Gli3*^*Δ699/Δ699*^ embryos (Fig. 2A) demonstrating that PDGFRα signalling was impaired in *Gli3*^*Δ699/Δ699*^ embryos. This finding confirms the data of our previous report where we found a decreased PDGFRα signalling in *Shh*^-/-^ mouse embryonic ventricles (Gerhardt et al., 2013). Both studies point to a scenario in which HH signalling positively regulates PDGFRα signalling in the ventricles of mouse embryos. In line with this scenario, *Pdgfrα* mutant, *Erk2* mutant and *Akt* mutant mouse embryos exhibit VSDs (Bax et al., 2010; Chang et al., 2010; Frémin et al., 2015; Schatteman et al., 1995). Both signalling pathways, HH signalling and PDGFRα signalling are mediated by primary cilia in mouse embryonic ventricles and, interestingly, ventricular cilia were shorter in *Gli3*^*Δ699/Δ699*^ embryos but not in *Gli3*^*XtJ/XtJ*^ embryos (Fig. 2B). Thus, the loss of GLI3 does not affect cilia length but it seems so as if the decreased HH signalling shortens cilia length. Based on this assumption, the question arises whether HH signalling regulates PDGFRα signalling directly, for example by controlling the expression of *Pdgfrα*, or indirectly, for example, by governing cilia length and function as ciliary length alterations are indicative for ciliary dysfunction (Cui et al., 2011; Dowdle et al., 2011; Garcia-Gonzalo et al., 2011; Larkins et al., 2011). Interestingly, previous reports show that GLI2 deficiency increases cilia length by reducing WNT signalling (shown in MEFs) and by enhancing autophagy (shown in NIH3T3 cells) (Hsiao et al., 2018; Rosengren et al., 2018). Here, we report that the length of ventricular cilia was unaltered in the absence of GLI3 but it was decreased in the presence of only GLI3-R (Fig. 2B). Consistent with the general assumption, namely that GLI2 acts as the predominant transcriptional activator and GLI3 as the predominant transcriptional repressor (Bai et al., 2002; Park et al., 2000; Wang et al., 2000), both GLI2 deficiency and the presence of GLI3-R alone caused reduced HH signalling (Fig. 1B, C) (Rosengren et al., 2018). Nevertheless, the consequences on cilia length are different. Potentially, the reason for this difference might be disparate cilia length control mechanisms in various cell types.

### Cell proliferation rate is decreased at ciliated areas of *Gli3*^*Δ699/Δ699*^ embryonic ventricles

In an earlier work, we observed a correlation between the proliferation of cells which are present at the ciliated regions of ventricles in mouse embryos and the outgrowth of the VS. In this context, we proposed a model indicating that cilia-mediated PDGFRα signalling regulates cell proliferation in distinct ventricular areas and that this proliferation drives the outgrowth of the VS (Gerhardt et al., 2013). Thus, we quantified proliferation in the ciliated regions of *Gli3*^*Δ699/Δ699*^ and *Gli3*^*XtJ/XtJ*^ embryonic ventricles by using phospho-histone H3 (pH3) and KI67 to mark proliferating cells. The proliferation rate was decreased in *Gli3*^*Δ699/Δ699*^ ventricles and unchanged in *Gli3*^*XtJ/XtJ*^ embryonic ventricles (Fig. 3A, B). These data confirm the previously observed correlation between the occurrence of VSDs and the reduced proliferation in ciliated areas of embryonic ventricles.

**Figure 3:**
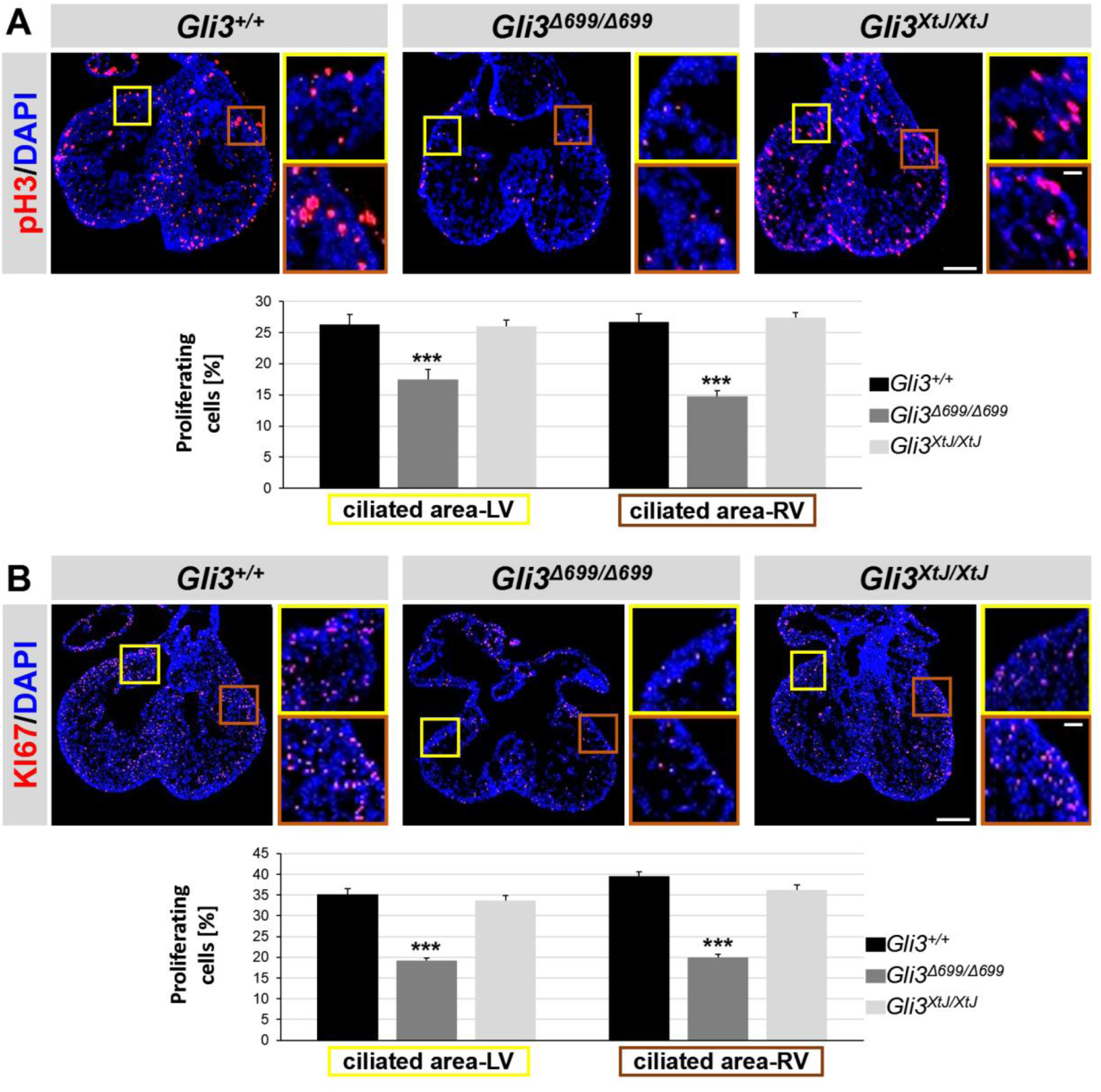
Reduced proliferation rate in ciliated regions of *Gli3*^*Δ699/Δ699*^ embryonic hearts. (A, B) Immunofluorescence on mouse embryonic heart sections obtained from WT (n=12), *Gli3*^*Δ699/Δ699*^ (n=8) and *Gli3*^*XtJ/XtJ*^ (n=12) embryos at E12.5. Proliferating cells are marked by pH3 (A) or KI67 (B). Cell nuclei are marked by DAPI. Scale bars represent a length of 100 µm in the overview and 15 µm in the insets. LV, left ventricle; RV, right ventricle.

### Rescue of PDGFRα signalling restores cilia-regulated proliferation in *Gli3*^*Δ699/Δ699*^ mutants

To examine whether this proliferation is indeed controlled by PDGFRα signalling, we turned to *in vitro* drug treatment experiments. We tried to isolate ventricular cells from the ciliated regions of wild-type and *Gli3*^*Δ699/Δ699*^ embryos but, unfortunately, all the ventricular cells stopped proliferation after the isolation. Next, we focussed on the analysis of *Gli3*^*Δ699/Δ699*^ mouse embryonic fibroblasts (MEFs). Firstly, we investigated PDGFRα signalling, cilia length and cell proliferation in order to check if *Gli3*^*Δ699/Δ699*^ MEFs show the same defects as *Gli3*^*Δ699/Δ699*^ ciliated ventricular cells. In these MEFs, the ciliary amount of PDGFRα, cilia length and proliferation rate were reduced (Fig. 4A-C) reflecting the defects of *Gli3*^*Δ699/Δ699*^ ciliated ventricular cells. Thus, the suggestion discussed above, namely that the cilia length differences observed between GLI2-deficient MEFs and *Gli3*^*Δ699/Δ699*^ embryonic ventricles could appear due to cell type-specific mechanisms underlying cilia length control, is questionable as these differences also occur in GLI2-deficient MEFs and *Gli3*^*Δ699/Δ699*^ MEFs. Consequently, the reason why cilia length is decreased in the presence of GLI3 alone remains elusive and should be addressed in future.

**Figure 4:**
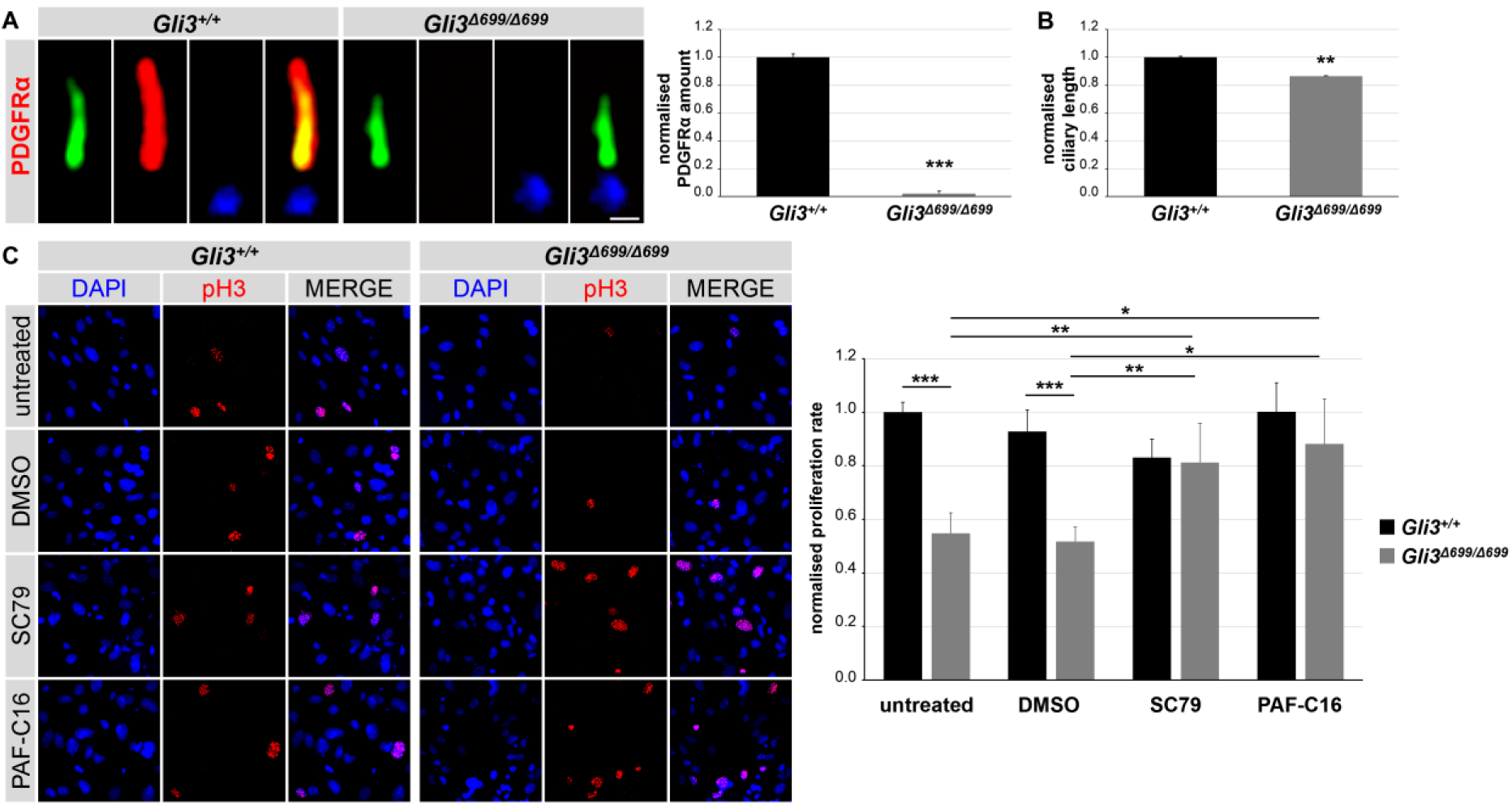
PDGFRα signalling activators rescue proliferation rate in *Gli3*^*Δ699/Δ699*^ MEFs. (A-C) Immunofluorescence on and ciliary length quantifications in MEFs obtained from WT and *Gli3*^*Δ699/Δ699*^ (n = 4, respectively) embryos at E12.5. (A) The ciliary axoneme is stained in green by acetylated α-tubulin, the basal body is stained in blue by γ-tubulin. The amount of PDGFRα is significantly reduced in cilia of *Gli3*^*Δ699/Δ699*^ MEFs. The scale bar represents a length of 0.5 µm. (B) Ciliary length measurement. (C) Proliferating cells are marked by pH3. Cell nuclei are marked by DAPI.

Since *Gli3*^*Δ699/Δ699*^ MEFs and *Gli3*^*Δ699/Δ699*^ embryonic ventricles show the same defects regarding PDGFRα signalling, cilia length and proliferation, we used *Gli3*^*Δ699/Δ699*^ MEFs for the drug treatment experiments. The AKT activator SC79 and the MEK1/2 activator PAF-C16 were administered to wild-type and *Gli3*^*Δ699/Δ699*^ MEFs. Upon treatment with SC79 as well as with PAF-C16, the proliferation rate of *Gli3*^*Δ699/Δ699*^ MEFs was rescued (Fig. 4C) demonstrating that the decreased PDGFRα signalling caused the reduced proliferation in *Gli3*^*Δ699/Δ699*^ MEFs.

The VSD is the most common congenital heart disorder in humans (Bjornard et al., 2013; Botto et al., 2001). Often, VSDs do not cause death in childhood but result in a severely impaired cardiac function in adulthood (Brickner et al., 2000; Srivastava, 2004). Today, the only curative treatment of VSDs is surgical repair. Mostly, surgical closure of VSDs is successful but it harbours several risks such as chronotropic incompetence, operative mortality, or late death (Heiberg et al., 2017; Schipper et al., 2017; Scully et al., 2010). For this reason, the development of alternative therapies against VSDs is important. Since the molecular causes of VSDs are largely unknown (Franco et al., 2006; Sakata et al., 2002), studies investigating these causes are urgently needed. Our data reveal how mutations in *Gli3* give rise to VSDs and help to pave the way for the development of therapies against VSDs suggesting that SC79 and PAF-C16 might be useful agents in VSD therapies.

The clinical relevance of our study is reflected by the fact that some mutations in *GLI3* also lead to VSDs in humans. Mutations in *GLI3* cause severe syndromes including the Greig cephalopolysyndactyly syndrome or the Pallister-Hall syndrome (Kang et al., 1997; Vortkamp et al., 1991). While the Greig cephalopolysyndactyly is caused by a complete loss of GLI3 function, patients suffering from the Pallister-Hall syndrome lack GLI3-A but have functioning GLI3-R (Biesecker, 1997; Johnston et al., 2005). Importantly, VSDs have never been described in patients suffering from the Greig cephalopolysyndactyly syndrome but Pallister-Hall syndrome patients develop VSDs (Démurger et al., 2015). Consequently, the human data resemble the results we obtained from mouse embryos and hence it is an exciting topic for future studies to evaluate whether our mouse data are transferable to humans.

## MATERIALS AND METHODS

### Antibodies

We used primary antibodies targeting actin (#A2066; Sigma-Aldrich), pAKT (#66444-1; Proteintech), GLI2 (#AF3635; R&D Systems), KI67 (#sc-7846; Santa Cruz Biotechnology, Inc.), PDGFRα (#sc-338; Santa Cruz Biotechnology, Inc.), pH3 (#06-570; EMD Millipore), pMEK1/2 (#9121; Cell Signaling Technology), acetylated α-tubulin (#sc-23950; Santa Cruz Biotechnology, Inc.), detyrosinated α-tubulin (#AB3201; EMD Millipore) and γ-tubulin (#sc-7396; Santa Cruz Biotechnology, Inc. and #T6557; Sigma-Aldrich).

### Cell culture and drug treatment

MEFs were isolated from single mouse embryos at embryonic day (E)12.5 after standard procedures. Cells were grown in DMEM containing 10% fetal calf serum (FCS), 1/100 (v/v) L-glutamine (Gibco), 1/100 (v/v) sodium pyruvate (Gibco), 1/100 (v/v) non-essential amino acids (Gibco) and 1/100 (v/v) pen/strep (Gibco) at 37°C and 5% CO_2_ until they reach confluency. Ciliogenesis was induced by serum starvation for at least 24 hours. Cells were treated with 10 µM SC79 (#SML0749; Sigma-Aldrich), 10 µM PAF-C16 (#sc-201009; Santa Cruz Biotechnology, Inc.) or DMSO (vehicle control) for 1 hour, respectively.

### Hematoxylin and eosin staining

Embryos at E14.5 were fixed in 4% PFA overnight at 4°C. Afterwards, they were gradually dehydrated using serial ethanol dilutions and finally embedded in paraffin. 12 μm sections were prepared. The sections were stained with haematoxylin and eosin (HE).

### Image acquisition

HE staining images were obtained at room temperature using the Zeiss Axioskop 2 system, an AxioCam MRc (Carl Zeiss AG) and the AxioVision Rel. 4.7.1 software package (Carl Zeiss AG). Fluorescence image acquisition and was performed at room temperature using a Zeiss Imager.A2 microscope, 100x, NA 1.46 oil immersion objective lens (Carl Zeiss AG) for ciliary protein measurements or a 20x, NA 0.5 objective lens (Carl Zeiss AG) for proliferation measurements. A monochrome charge-coupled device camera (AxioCam MRm, Carl Zeiss AG), and the AxioVision Rel. 4.8 software package (Carl Zeiss AG) were used. Appropriate anti-mouse, -rabbit and -goat Dylight405, Dylight488, Cy3, Alexa405 and Alexa488 antibodies were used as fluorochromes.

### Immunofluorescence

Mouse embryos at E12.5 were isolated and embryonic hearts were dissected. Hearts were fixed in 4% PFA for 1 hour and incubated in 30% sucrose (in PBS) overnight at 4°C. At the next day, embryonic hearts were embedded in Tissue-Tec O.C.T. compound (#4583; Sakura Finetechnical) and freezed at −80°C. Transverse cryostat sections (6 µm) were prepared. Heart sections were rinsed in PBS and permeabilised with PBS/0.5% Triton X-100 for 10 min. After three washing steps with PBS/0.1% Triton X-100 blocking was performed with 10% donkey serum in PBST. Subsequently the sections were incubated with primary antibodies diluted in PBS/0.1% Triton X-100 overnight at 4°C. On the next day, the sections were washed three times with PBS/0.1% Triton X-100 and incubated with the secondary antibodies diluted in PBS/0.1% Triton X-100 for 2 hours. After several washing steps the sections were embedded in Mowiol optionally containing DAPI (#1.24653; Merck).

For immunofluorescence on MEFs, cells were plated on coverslips until the reach confluency. MEFs were serum-starved for at least 24 hours. Cells were fixed with 4% PFA for 1 hour at 4°C. Fixed MEFs were rinsed three times with PBS, followed by a permeabilisation step with PBS/0.5% Triton-X-100 for 10 minutes. The samples were washed three times with PBS and blocking was performed by incubation in PBST (PBS/0.1% Triton-X-100) containing 10% donkey serum for at least 30 minutes at room temperature. Diluted primary antibodies (in PBS/0.1% Triton-X-100) were incubated overnight at 4°C. After three washing steps with PBST, incubation with fluorescent secondary antibody (in PBS/0.1% Triton-X-100) was performed at room temperature for 1 hour followed by several washing steps and subsequent embedding with Mowiol optionally containing DAPI (#1.24653; Merck).

### Statistical analysis

Data are presented as mean ± standard error of mean (SEM) and statistics for all data were performed using Prism (GraphPad). Two-tailed *t-*tests with Welch’s corrections were performed for all data in which two datasets were compared. Analysis of variance (ANOVA) and Tukey honest significance difference (HSD) tests were used for all data in which more than two datasets were compared. A P-value <0.05 was considered to be statistically significant (one asterisk), a P-value <0.01 was defined as statistically very significant (two asterisks) and a P-value <0.001 was noted as statistically high significant (three asterisks).

### qRT-PCR

We isolated RNA from single mouse embryonic hearts at embryonic stage E12.5 by using the RNeasy Kit (#74104; Qiagen) and the RNase-Free DNase Set (#79254; Qiagen). The isolated heart RNA was converted into cDNA by QuantiTect Reverse Transcription Kit (#205311; Qiagen). Quantitative real-time PCR was performed using a Step One Real-Time PCR System Thermal Cycling Block (#4376357; Applied Biosystems) and a Maxima SYBR Green/ROX qPCR Master Mix 2x (#K0222; Thermo Scientific). The following primer sets were used: *Gli1* (forward 5’-TACCATGAGCCCTTCTTTAG-3’; reverse 5’-TCATATCCAGCTCTGACTTC-3’), *Ptch1* (forward 5’-CAACCAAACCTCTTGATGTG-3’; reverse 5’-CCTGCCAATGCATATACTTC-3’), *Hprt* (forward 5’-AGGGATTTGAATCACGTTTG-3’; reverse 5’-TTTACTGGCAACATCAACAG-3’).

We used 50 ng of embryonic cDNA of each sample in triplicate reactions for the real-time PCR. The sample volume of 25 µl contains 100 nM primers and 50 nM probe. Cycling conditions were 50°C for 2 min and 95°C for 10 min, followed by a 40-cycle amplification of 95°C for 15 s, 60°C for 30 s and 72°C for 30s. The analysis of real-time data was performed by using included StepOne Software version 2.0.

### Quantifications

Quantification of ciliary protein amounts and proliferation rates were realised by using Image J (National Institutes of Health).

Intensity measurement of ciliary proteins (PDGFRα, pAKT and pMEK1/2) based on immunofluorescence staining was performed as described before (Garcia-Gonzalo et al., 2011; Garcia-Gonzalo et al., 2015; Gerhardt et al., 2015; Roberson et al., 2015; Struchtrup et al., 2018; Wiegering et al., 2018; Yee et al., 2015). The ciliary length has to be taken into account while quantifying the ciliary amount of PDGFRα, pAKT and pMEK1/2 in different genotypes. Therefore we used the area marked by acetylated α-tubulin as a reference and quantified the average pixel intensity of the PDGFRα, pAKT and pMEK1/2 staining. To exclude unspecific staining from the measurements, we subtracted the mean value of the average pixel intensity of three neighboring regions free from specific staining.

Proliferation quantifications were performed via cell counting. Therefore, we investigated six sections for each experimental embryo (heart sections) or three different regions of two coverslips per individual originated MEFs. Total cell number was counted via DAPI. Proliferating cells were counted via KI67- or pH3-positive signals and related to the total cell number. Finally, the proliferation rates were normalized to WT.

### Western Blotting

Western blot studies were performed as described earlier using an anti-GLI2 (#AF3635; R&D Systems) antibody (Wang et al., 2000). Anti-actin antibody (#A2066; Sigma-Aldrich) was used as loading control. Visualization of protein bands was realised by the LAS-4000 mini (Fujifilm). Band intensities were measured by using ImageJ (National Institutes of Health).

## ACKNOWLEDGEMENTS

The authors thank Matias Zurbriggen and Leonie-Alexa Koch for their generous help to enable the continuation of the study.

